# Non-pathogenic leaf-colonising bacteria elicit pathogen-like responses in a colonisation density-dependent manner

**DOI:** 10.1101/2023.05.04.539505

**Authors:** Moritz Miebach, Léa Faivre, Daniel Schubert, Paula Jameson, Mitja Remus-Emsermann

## Abstract

Leaves are colonised by a complex mix of microbes, termed the leaf microbiota. Even though the leaf microbiota is increasingly recognised as an integral part of plant life and health, our understanding of its interactions with the plant host is still limited. Here, mature, axenically grown *Arabidopsis thaliana* plants were spray-inoculated with six diverse leaf-colonising bacteria. The transcriptomic changes in leaves were tracked over time and significant changes in ethylene marker (*ARL2*) expression were observed only two to four days after spray-inoculation. Whole transcriptome sequencing revealed that four days after inoculation, leaf transcriptional changes to colonisation by non-pathogenic and pathogenic bacteria differed in strength but not in the type of response. Inoculation of plants with different densities of the non-pathogenic bacterium *Williamsia* sp. Leaf354 showed that high bacterial titers caused disease phenotypes and led to severe transcriptional reprogramming with a strong focus on plant defence. An *in silico* epigenetic analysis of the data was congruent with the transcriptomic analysis. These findings suggest (1) that plant responses are not rapid after spray-inoculation, (2) that plant responses only differ in strength and (3) that plants respond to high titers of non-pathogenic bacteria with pathogen-like responses.

**Plain Language Summary:** Plants are colonised by diverse bacteria affecting many aspects of plant life. Here we show that plants do not differentiate between different bacteria but measure their quantities to keep bacterial numbers in check.

## Introduction

Plants are colonised by a vast variety of bacteria with various effects on plant health and growth, ranging from pathogens to nitrogen-fixing rhizobacteria (Wang *et al*., 2018; Kelly *et al*., 2018). Past research mainly focused on a few important microbiota members, overlooking most of the remarkably diverse microbes present on and within plants. Increasing evidence showcases the positive effects of the plant microbiota on its host, including the protection against abiotic (Lau & Lennon, 2012) and biotic (Innerebner *et al*., 2011; Ritpitakphong *et al*., 2016; Zengerer *et al*., 2018; Vogel *et al*., 2021) stressors, the promotion of growth (Spaepen *et al*., 2009) and the assimilation of specific nutrients (Hacquard *et al*., 2015; Singh *et al*., 2022). Some of these beneficial effects are solely attributed to microbes, whereas others involve the plant as an interaction partner. For example, with regard to the protection against biotic stress, some microbes can protect the plant from a pathogen via resource competition with the pathogen (Ji & Wilson, 2002) or the production of antimicrobials targeting the pathogen (Zengerer *et al*., 2018), whereas others stimulate the plant immune system, leading to increased immune responses upon subsequent infection with a pathogen (Pieterse *et al*., 1996; Vogel *et al*., 2016, 2021).

Even though the microbiota seems to be an integral part of plant life, our understanding of its interaction with the host is still limited. Studies on plant pathogens have shown that early perception of microbes is conferred by the detection of microbe-associated molecular patterns (MAMPs). MAMPs are conserved traits of microbes irrespective of their symbiotic relationship with the plant, raising the question if and how plants discriminate between pathogenic and beneficial bacteria. Recently, it was shown that plants respond to diverse non-pathogenic leaf colonisers, at varying intensities of transcriptional reprogramming (Maier *et al*., 2021). However, whether the amount of transcriptional reprogramming is caused by the different identities of the bacteria is unclear. Notably, the observed plant response was enriched for plant defence-associated genes and measured nine days after inoculation of ten-days-old seedlings, indicating persistently active immune responses. This is intriguing, since plant immune responses infer a growth penalty, commonly referred to as the growth-defence tradeoff (Huot *et al*., 2014; He *et al*., 2022).

As a persistent immune activation infers a growth penalty, plants need to activate their immune system according to potential pathogen threat in a timely manner (Huot *et al*., 2014; He *et al*., 2022). Different activation kinetics likely lead to different downstream signalling events and finally to a different immune response, highlighting the importance of time-resolved analyses. In other words, MAMPs are present early in the plant-microbe interaction and lead to early plant responses. However, MAMPs are less indicative of the symbiotic relationship between the microbe and the plant compared to effector proteins. Accordingly, plant responses triggered by MAMPs are more transient than those triggered by effector molecules (Gao et al., 2013; Lamb & Dixon, 1997; Tsuda et al., 2013). Plant responses to different MAMPs are almost identical early after MAMP application, but differ later, resulting in differential immune outputs (Zipfel *et al*., 2006; Kim *et al*., 2014; Bjornson *et al*., 2021). Elf18 and chitosan, for example, predominantly activate jasmonic acid (JA)-mediated immune responses early on, resulting in JA-mediated immunity later, while flg22 activates jasmonic acid (JA)-mediated and ethylene (ET)-mediated immune responses early on, resulting in salicylic acid (SA)-mediated immunity later (Kim *et al*., 2014). This further highlights the importance of time-resolved analyses and suggests that, already at the level of MAMP recognition, plant immune responses differ depending on the cocktail of MAMPs present.

The response of plants to leaf colonisation, whether by pathogenic or non-pathogenic bacteria, involves a significant transcriptome reprogramming (Moore *et al*., 2011; Maier *et al*., 2021). This reprogramming relies on the interaction of numerous transcription factors with the local chromatin environment. Indeed, several histone modifications such as H3K4me3, H3K36me3 and lysine acetylation were found to contribute to the induction of genes following pathogen exposure (Berr *et al*., 2012; Ding & Wang, 2015). By contrast, transcriptional reprogramming following colonisation by non-pathogenic bacteria has not been investigated.

To monitor the dynamics of plant immune responses, we inoculated plants with diverse bacteria, including a plant pathogen, and measured transcriptional outputs, representing the three major phytohormone pathways in plant immunity (SA, JA and ET), at various times after inoculation. As the strongest responses were observed 96 hours post inoculation (hpi), we measured the whole transcriptional plant responses at this time. Interestingly, plant responses to various bacterial inoculants strongly overlapped and we observed a trend that the plant responses were dependent on bacterial density. Most differentially expressed genes in response to one isolate were also differentially expressed in response to isolates that exhibited overall stronger responses. The question then arose whether non-pathogenic bacteria could induce plant defence responses at artificially high colonisation densities. To address this question, we assessed the effects of different bacterial densities of the non-pathogenic leaf coloniser *Williamsia* sp. Leaf354 on *in planta* transcription as well as on plant health and plant weight 96 hpi and 21 dpi, respectively, to test the hypothesis that plants are monitoring bacterial population density rather than differentiating between different bacterial colonisers. Finally, in an effort to uncover chromatin marks that might contribute to transcriptional responses to non-pathogenic leaf colonisers, we performed an *in silico* chromatin analysis.

## Materials and Methods

### Plant growth

Plants were grown as previously described in (Miebach *et al*., 2020). Briefly, sterilised seeds were germinated on ½ MS (Murashige and Skoog medium, including vitamins, Duchefa, Haarlem, Netherlands) 1% phytoagar (Duchefa) filled pipette tips.

Healthy looking seedlings were aseptically transferred, without removal from the pipette tip, aseptically into Magenta boxes (Magenta vessel GA-7, Magenta LLC, Lockport, IL, USA) filled with ground Zeolite (sourced from cat litter - Vitapet, Purrfit Clay Litter, Masterpet New Zealand, Lower Hutt, New Zealand) and watered with 60 mL ½ MS. Each box received four seedlings. The boxes were closed with lids that allowed for gas exchange and placed into a climate cabinet (85% relative humidity, 11 h light, 13 dark, 21 °C, 150-200 µmol light intensity). Plants were grown for four weeks for the time course and six weeks for the bacterial density experiment before they were treated with bacteria or mock controls.

### Plant inoculation

Bacterial suspensions were prepared as previously described in (Miebach *et al*., 2020). Briefly, bacteria were cultivated at 30 °C on minimal media agar plates containing 0.1% pyruvate as a carbon source. Bacterial suspensions were prepared from bacterial colonies suspended in phosphate-buffered saline (PBS, 0.2 g L^−1^ NaCl, 1.44 g L^−1^ Na_2_HPO_4_ and 0.24 g L^−1^ KH_2_PO_4_) and washed twice via centrifugation at 4000 × *g* for 5 min followed by discarding the supernatant and again adding PBS. Table **1** contains the list of bacteria used in this study. For the time course experiment the optical density (OD_600 nm_) was adjusted so that the suspension contained 2 × 10^7^ colony forming units (CFU) ml^-1^. To explore the influence of bacterial load on plant responses, the bacterial suspensions were adjusted to 10^5^, 10^6^, 10^7^ and 10^8^ CFU ml^-1^. Next, 200 µL (time course experiment) or 1 ml (bacterial load experiment) of bacterial solution was sprayed per plant tissue culture box using an airbrush spray gun (0.2 mm nozzle diameter, Pro Dual Action 3 #83406). To obtain a homogeneous coverage, the distance between the airbrush spray gun and the plants was increased by stacking a plant tissue culture box, with the bottom cut off, onto the plant tissue culture box containing the plants being spray-inoculated.

**Table 1:** List of bacterial strains used in this study.

| Bacterial Strain | Reference | Abbreviation |
| --- | --- | --- |
| *Pseudomonas syringae*<br>DC3000 | (Cuppels, 1986) | Pst |
| *Sphingomonas* sp. Leaf34 | (Bai *et al.*, 2015) | Sphingo34 |
| *Acidovorax* sp. Leaf84 | (Bai *et al.*, 2015) | Acido84 |
| *Microbacterium* sp. Leaf347 | (Bai *et al.*, 2015) | Micro347 |
| *Williamsia* sp. Leaf354 | (Bai *et al.*, 2015) | Willi354 |
| *Pedobacter* sp. Leaf194 | (Bai *et al.*, 2015) | Pedo194 |

### Bacterial enumeration

Above ground plant parts were detached from belowground parts using sterilised equipment and placed individually into pre-weighed 1.5 mL tubes. After determining the plant weight 1 mL PBS 0.02% Silwet L-77 (Helena Chemical Company) was added to each tube. Bacteria were dislodged from the sample by shaking twice at 2.6 m s^-1^ for 5 min (Omni Bead Ruptor 24) and sonication for 5 min. Bacterial CFU were enumerated using plate counting on R2A media plates.

### Gene expression analysis

Four-weeks-old and six-weeks-old plants were spray-inoculated with individual strains (Table **1**) or PBS (mock control) for the time course and bacterial load experiment, respectively. For the time course experiment above ground plant parts were harvested after 1, 3, 6, 9, 12, 24, 48, and 96 h post inoculation (hpi). For the bacterial load experiment above ground plant parts were harvested 96 hpi. The plant material was collected in RNase-free microcentrifuge tubes (MCT-150-C, Axygen, Corning, USA) and was then immediately flash frozen in liquid N_2_. Two plants from different growth boxes were pooled per tube to form a biological replicate. Three biological replicates were sampled per treatment and time point. Flash-frozen samples were ground to a fine powder in the collection tube using Teflon pestles (General Lab Supply, Lab Supply, Dunedin, New Zealand). RNA extraction was performed using the Isolate II RNA Plant kit (Bioline, London, England).

### RT-qPCR analysis

For cDNA synthesis, 1 µg of RNA was used and for the no Reverse Transcriptase (noRT) control, using the VitaScript First strand cDNA synthesis kit (Procomcure Biotech, Thalgau, Austria). RT-qPCR was performed using the 2× ProPlant SYBR Mix (Procomcure Biotech) in 15 µL reaction volumes with 0.2 µM of each primer and 0.001 g L^−1^ of initial RNA in the cDNA mix. QPCRs were run using the recommended protocol for 2× ProPlant SYBR Mix (Procomcure Biotech) on a Rotor-Gene Q (Qiagen, Hilden, Germany). Technical triplicates were performed for each sample. The ROX dye, present in the 2× ProPlant SYBR Mix, was used to normalise for master mix variation between tubes. A mix of equal amounts of all cDNAs was used for normalisation between runs. mRNA concentrations were calculated using Equation (1).

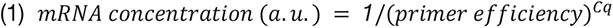

Primers that were first used in this study were designed using ‘primer-blast’ (NCBI, Bethesda, MD, USA). Primer efficiencies were determined via serial template dilutions (Nolan *et al*., 2013). The mRNA concentration of each target gene was then normalised against the mean mRNA concentration of two stably expressed, previously described reference genes (Table **2**, (Czechowski *et al*., 2005)). Next, the normalised mRNA concentration of each treatment (bacterial inoculation) was normalised against the mean normalised mRNA concentration of mock-treated samples to emphasise treatment-related changes in gene expression (Denoux *et al*., 2008).

**Table 2:**
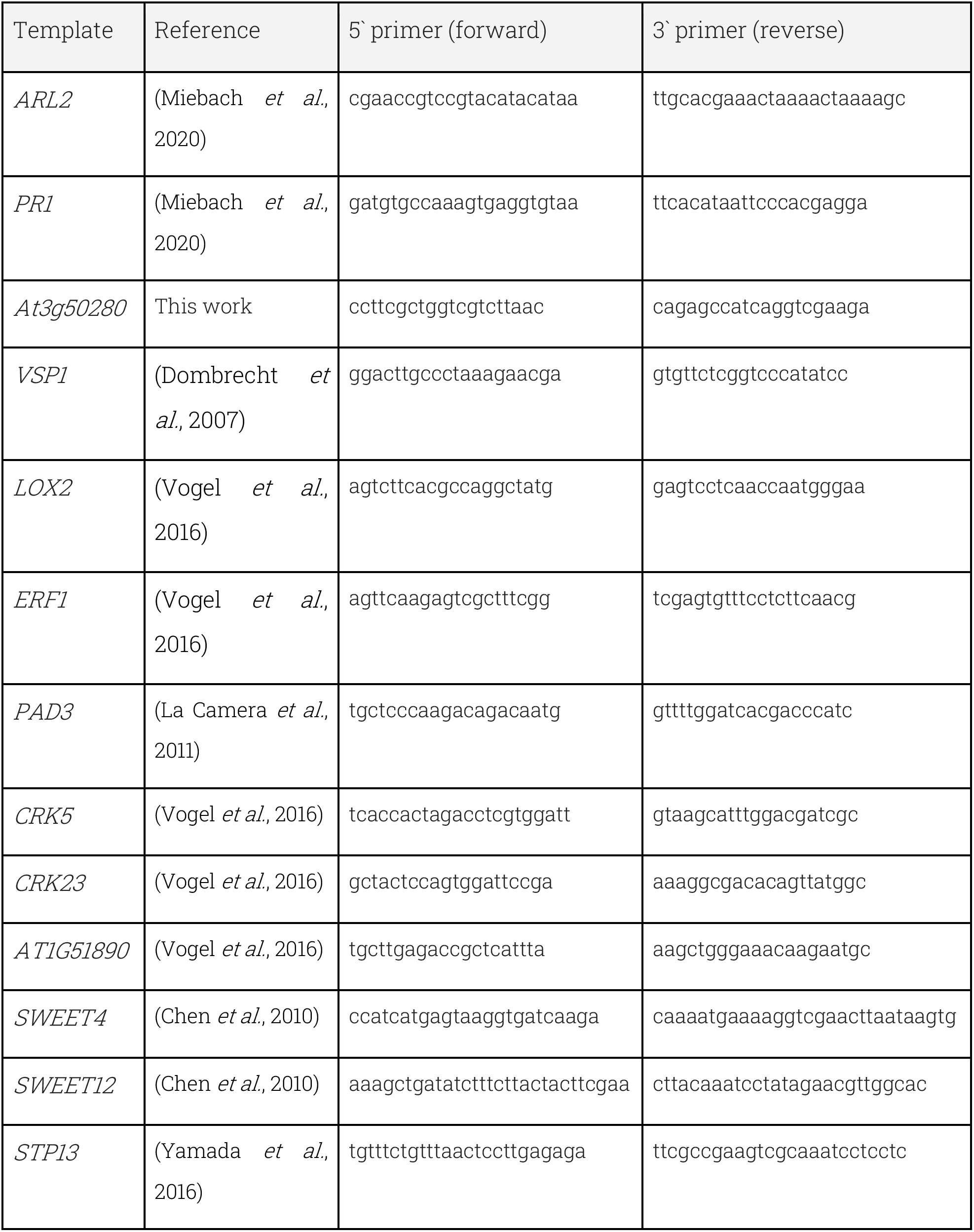
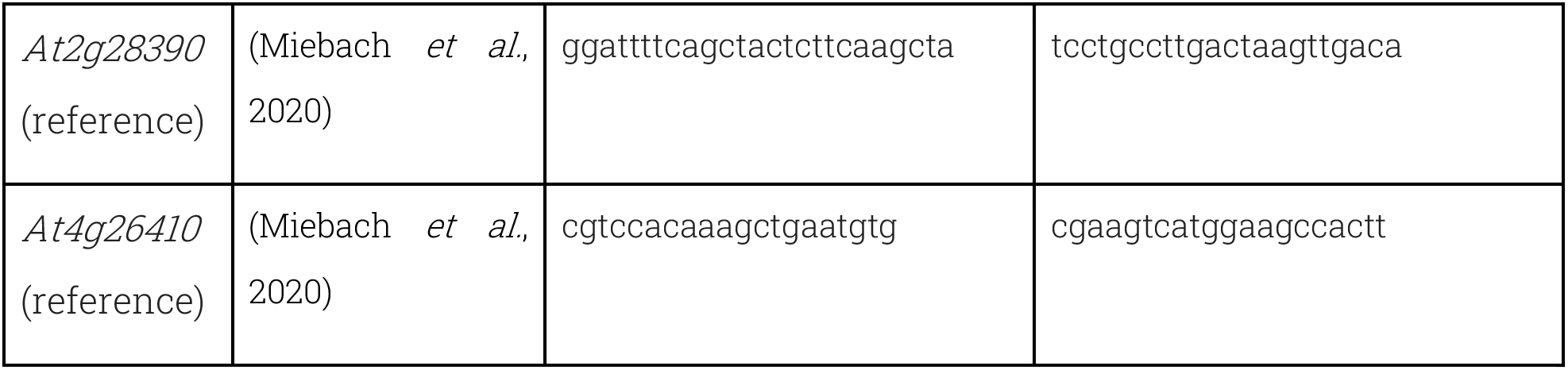
List of primers used in this study.

### RNA sequencing

RNA library preparation and sequencing was performed by Custom Science (Epping, Australia). RNA quality and purity was assessed using a Nanodrop (ND-2000 Spectrophotometer, Thermo Scientific, Waltham, MA, USA). RNA integrity was verified on a 2100 Bioanalyzer (Agilent, Santa Clara, CA, USA) using the RNA 6000 Nano Kit (Agilent). Poly-A enriched libraries were prepared using the NEBNext UltraTM RNA Library Prep Kit (New England Biolabs, Ipswich, USA) for Illumina. RNA sequencing was performed on a NovaSeq 6000 (Illumina, San Diego, USA) platform. Approximately 30,000,000 paired-end reads with a length of 150 bp were generated per sample. Adapters and low-quality reads (end sequences with base quality < 20 and sequences with N content > 10%) were removed from raw sequences and sequences below 75 bp were filtered using cutadapt (v1.9.1) (Martin, 2011). The resulting reads were mapped against the *Arabidopsis thaliana* (arabidopsis) reference genome (TAIR10). Mapped reads were counted with featureCounts (v1.22.2) (Liao *et al*., 2014). RNA sequencing data are available from the GEO repository under accession number GSE232254.

### RNA sequencing data analysis

Genes above 0.5 counts per million in at least three samples were used for differential gene expression analysis using edgeR (Robinson *et al*., 2010). Counts were scaled to effective library sizes by the trimmed mean of M values method (Robinson & Oshlack, 2010). Gene-wise dispersions were estimated via Cox-Reid profile-adjusted likelihood and squeezed to trended dispersions using an empirical Bayes method (McCarthy *et al*., 2012). Genes were determined as differentially expressed using the TREAT method under edgeRs general linear model framework (McCarthy & Smyth, 2009). Genes with a fold change (FC) significantly above log_2_(1.3) and below a False Discovery Rate (FDR; Benjamini-Hochberg *p* correction) cut-off of 5% were kept as differentially expressed genes (DEGs). The FC threshold was determined using elbow plots (Fig. **S3a**, Fig. **S4a**). MDS plots were generated from trimmed mean of M values method normalised gene counts. K-means clusters were calculated from log_2_-transformed counts per million, that were centred around the mean for each gene. A prior count of two was added to each gene count to prevent taking the logarithm of zero. Elbow plots were used to determine the optimal number of Ks (Fig. **S3b**, Fig. **S4b**). Heatmaps were generated from log_2_-transformed counts per million (cpm), that were centred around the mean for each gene. GO term enrichment analysis was performed using the PANTHER classification system (v17.0) (Mi *et al*., 2021).

### Chromatin state analysis

Chromatin state coordinates were obtained from (Sequeira-Mendes *et al*., 2014) and gene coordinates were obtained from the TAIR10 annotation from BioMart, Plantsmart28 (Durinck *et al*., 2005). A gene was considered to be in a certain state if at least 150 bp (approximate length of DNA wrapped around one nucleosome) of its gene body overlapped with the respective state. Thus, a single gene may have several distinct states along its coding sequence. Genes induced or repressed after Pst exposure were divided into two sets depending on whether they were also differentially expressed by exposure to Micro347 or Willi354 (Non Pst-specific) or only by Pst (Pst-specific). Similarly, genes induced or repressed by the highest inoculation density of Willi354 were divided into two sets depending on whether they were also differentially expressed by exposure to lower bacterial densities (non 10^8^-specific) or not (10^8^-specific). The proportion of genes of interest in each chromatin state was compared with the proportion of genes in the respective state in the complete genome. The significance of the difference between the two proportions was tested using the Marascuilo procedure, with a confidence level of 0.95.

## Results

The aim of this study was to broaden our knowledge of the intricate relationship between the plant and its bacterial colonisers. The focus lay on assessing plant transcriptional responses to a diverse array of microbial colonisers. Therefore, six microbial leaf colonisers representing all major phyla of the core leaf microbiota were selected (Vorholt, 2012; Bai *et al*., 2015) (Fig. **1**).

**Fig. 1:**
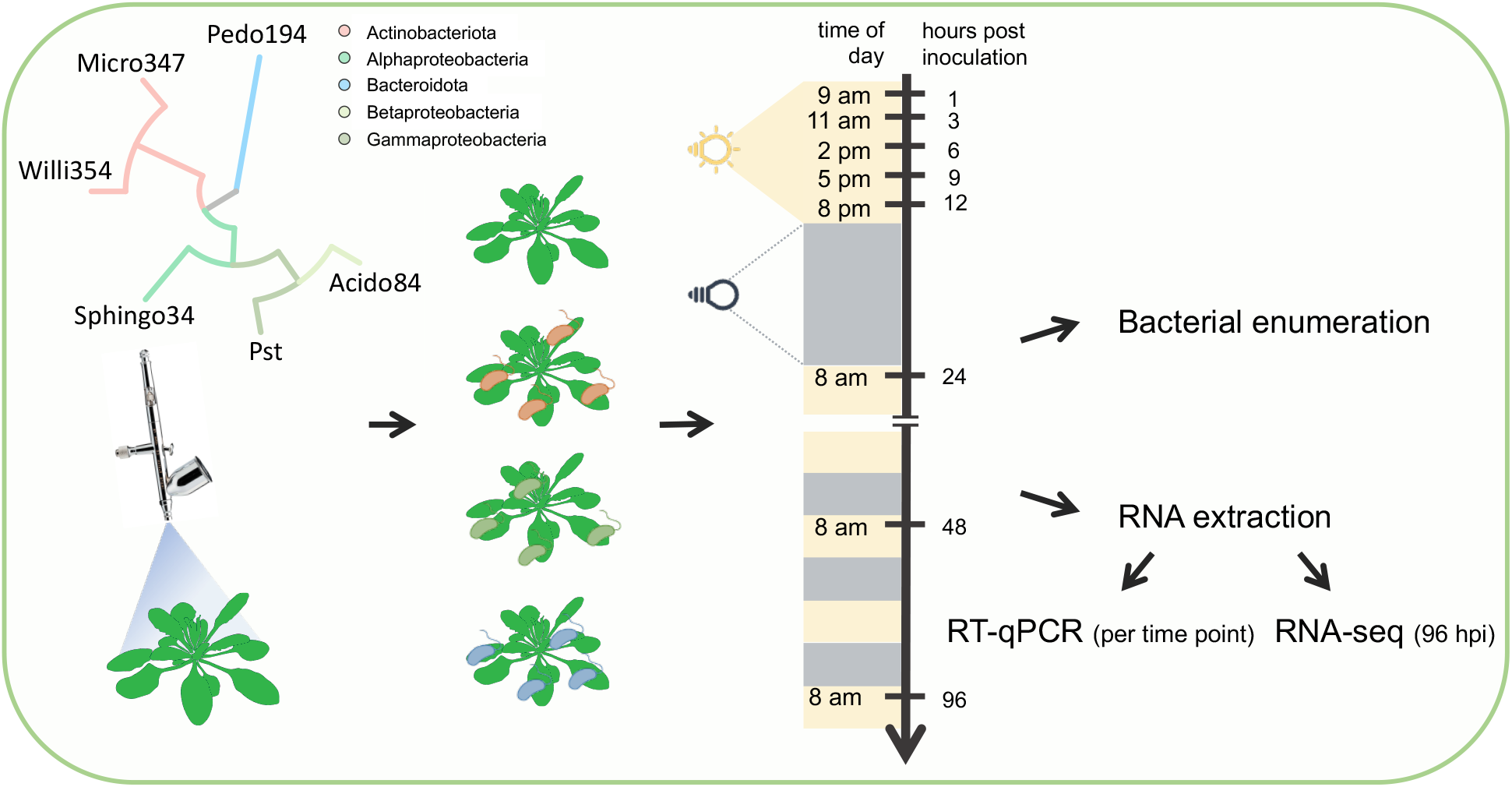
Experimental design. Four-weeks-old axenically-grown *Arabidopsis thaliana* plants were spray-inoculated with individual bacterial strains, depicted in the phylogenetic tree in the top left corner of the figure. Plants were harvested at different times after inoculation. Some plants were used for bacterial enumeration and others for RNA extraction.

### Temporal responses of the plant immune system

Arabidopsis plants were grown axenically in the ‘Litterbox’ system, to ensure (1) low artificial, but strictly controlled, growth conditions and (2) to prevent a strong inoculation of the growth media post inoculation (Miebach *et al*., 2020). Four-weeks-old plants were spray-inoculated with individual strains of bacterial leaf colonisers, at ∼ 10^5^ - 10^6^ bacteria per g of leaf, the bacterial carrying capacity of plants in temperate environments (Kniskern *et al*., 2007; Reisberg *et al*., 2012; Rastogi *et al*., 2012; Burch *et al*., 2016; Gekenidis *et al*., 2017) . The temporal course of bacterial densities post inoculation confirmed that bacteria were sprayed close to carrying capacity. Within 4 dpi the bacterial densities for Sphingo34 and Micro347 remained stable at 10^6^ bacteria per g of leaf. In contrast, the bacterial densities for Willi354 and Pst slightly rose to 10^7^ bacteria per g of leaf within 4 dpi (Fig. **2a**). Interestingly, the bacterial density of Micro347 significantly (*p* < 0.001, Tukey’s HSD test) dropped to ∼ 10^4^ bacteria per g of leaf 7 dpi. This two-magnitude drop in bacterial density cannot be explained by the increase in plant weight (Fig. **S1**) and, therefore, suggests that the bacteria were dying.

**Fig. 2:**
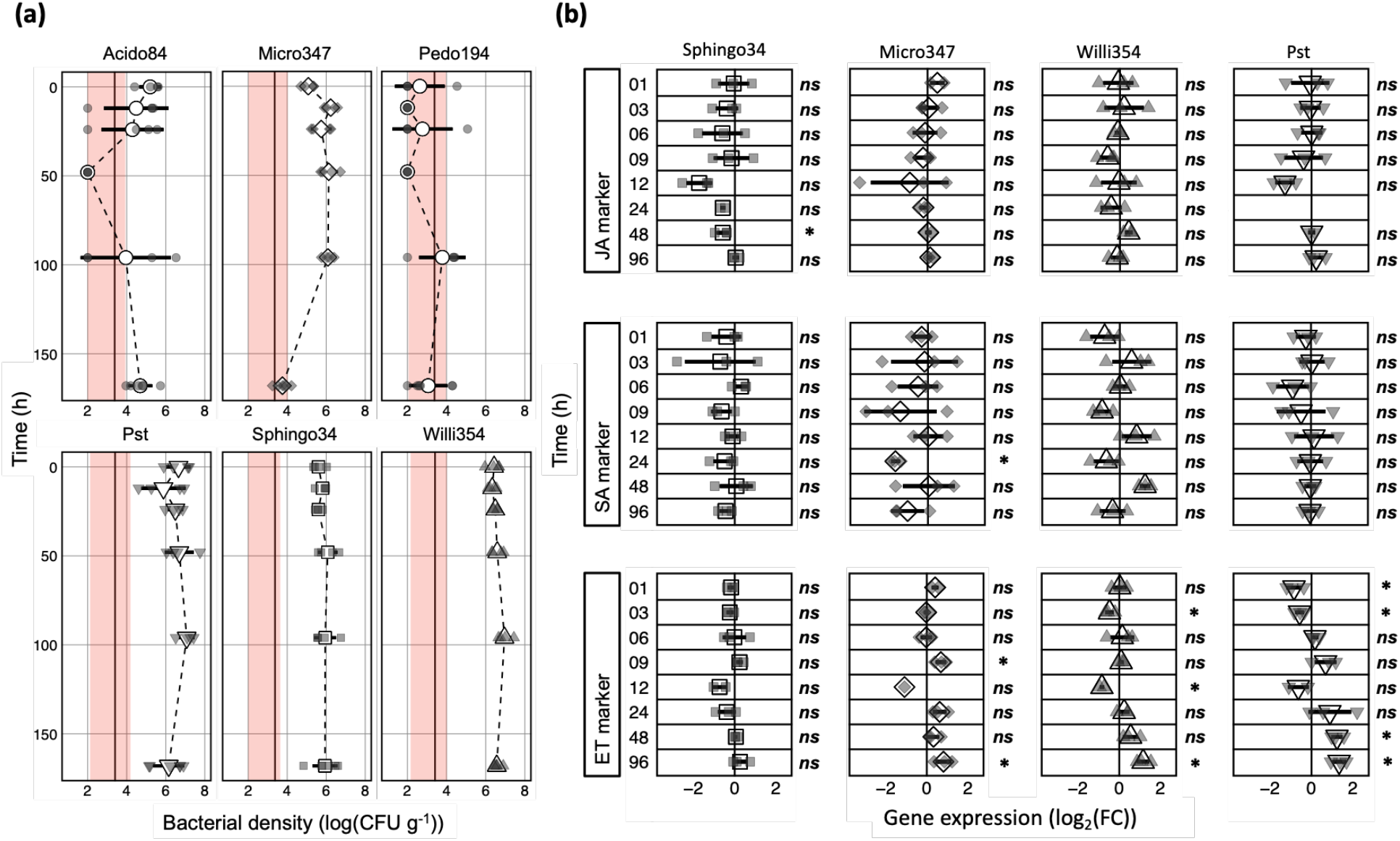
Temporal transcriptional responses to bacterial colonisation. Four-weeks-old axenically-grown arabidopsis plants were spray-inoculated with individual bacterial strains (see coloured boxes; strains are sorted alphabetically in (**a**) and by response amplitude of the ethylene marker gene in (**b**)). (a) Bacterial density on above ground plant parts at several times post inoculation. To accommodate plants with CFU below the limit of detection, 10^2^ CFU g^-1^ were added to the count of every sample. (**b**) Depicted are log_2_(FC) in gene expression relative to mock-treated control for a jasmonic acid (JA) marker gene (*At3g50280*), a salicylic acid (SA) marker gene (*PR1*) and an ethylene (ET) marker gene (*ARL2*). Large white shapes depict the mean, bars depict standard deviation, smaller grey shapes depict individual biological replicates, dashed black line connects the means, solid black line represents the lower limit of detection based on mean plant weight and the red areas represent the range of the limit of detection based on the heaviest and lightest plant, * depicts statistical significance (Bonferroni corrected *p* < 0.05) and *ns* depicts the lack of statistical significance, T-test.

Two out of the six bacterial leaf colonisers, Acido84 and Pedo194, failed to consistently establish densities above the threshold of detection. Acido84 was recovered from some, but not all plants. Whenever Acido84 was recovered it reached densities of ∼ 10^5^ - 10^6^ CFU g^-1^. This heterogeneity in colonisation success was unlikely to have been caused by non-homogeneous spray inoculation, as other inoculants exhibited considerably lower plant to plant variation (Fig. **2a**). Further, all plants sampled 168 hpi harboured ∼ 10^4^ - 10^6^ CFU g^-1^.

Plants inoculated by one of the four successful coloniser strains, Micro347, Pst, Sphingo34 and Willi354, were investigated further by qPCR (Fig. **2b**). Early temporal changes in the plant immune response were tracked using previously reported marker genes that follow the levels of the three major phytohormones in plant immunity: ET, JA and SA (Kim *et al*., 2014). Marker gene expression was measured at eight different time points ranging from 1 hpi to 96 hpi. As dynamic changes in expression were expected early after inoculation, five of the eight measurement times fell within the first 12 hpi.

Surprisingly, gene expression changes caused by the bacterial treatments were relatively weak. The strongest changes did not exceed 5-fold (maximum mean: 2.5-fold; maximum individual replicate: 4.6-fold) in gene expression, relative to the mock-treated control (Fig. **2b**). The strongest changes were observed in the expression of the ET marker, *ARL2*. Early after inoculation, its expression dropped significantly, at 1 and 3 hpi for Pst and 3 hpi for Willi354. After recovering to the expression levels found in mock-treated plants the relative expression of the ET marker dropped again at 12 hpi, which was significant in the case of Willi354, with Pst seemingly following the same trend. Expression levels then rose above mock-treated control by ∼ 2.5-fold at 96 hpi, with changes being statistically significant for 48 and 96 hpi for Pst and 96 hpi for Willi354 (Fig. **2b**). In addition, Micro347 showed a significant ∼ 1.8-fold rise in ET marker gene expression at 96 hpi (Fig. **2b**).

Regarding the JA marker, statistically significant changes were only observed in response to Sphingo34. A drop in expression was observed between 12 and 48 hpi with the latter being statistically significant, but rather weak at ∼ 1.5-fold (Fig. **2b**). Expression of the SA marker, *PR1*, fluctuated strongly, but not significantly between up- and downregulation upon Willi354 treatment. Strong fluctuations between no expression change and a strong downregulation were observed following Micro347 treatment with a significant ∼ 3-fold decrease in expression at 24 hpi. Sphingo34 and Pst elicited no changes in *PR1* expression (Fig. **2b**).

The increase in ET marker expression within the last three sampling times (24, 48, 96 hpi) seemed to follow the increase in bacterial density of the inoculants (Fig. **2a, b**).

Interestingly, 57% (adj. R^2^ = 0.57, *p* = 0.0027) of the relative to the mock control can be explained by change in gene expression the bacterial density, irrespective of the inoculant (Fig. **3**).

**Fig. 3:**
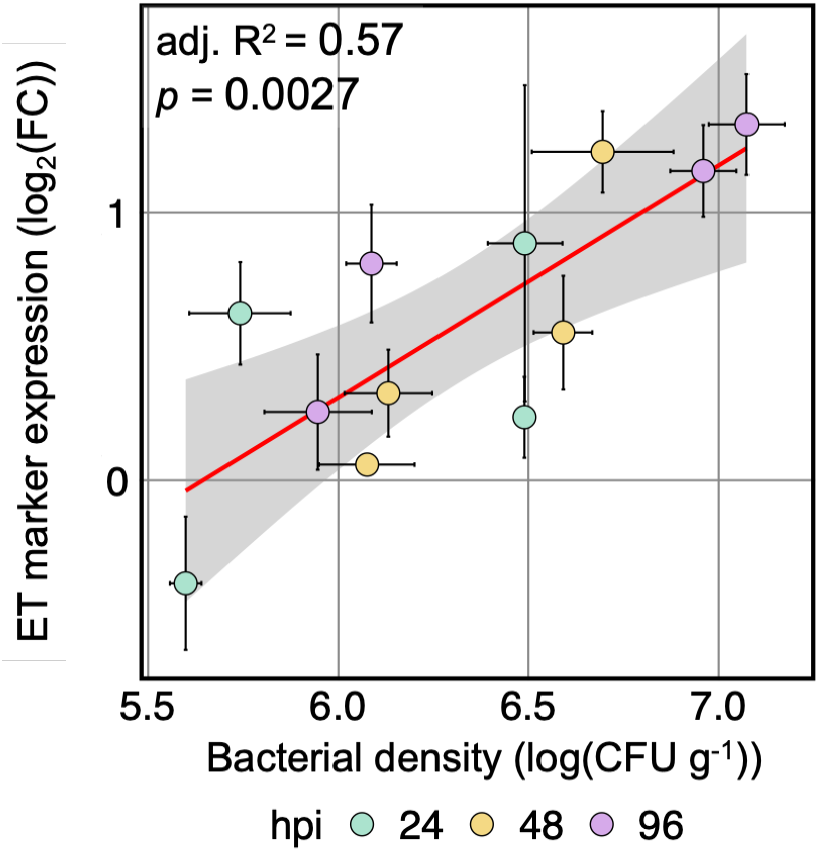
Ethylene marker responses may be explained by bacterial density. Correlation between mean log_2_(FC) in gene expression of ethylene (ET) marker gene (*ARL2*) relative to mock-treated control and mean bacterial density. Colours depict the time of sampling post inoculation, red line depicts a fitted linear model, the grey bar depicts limits of 95% confidence interval, error bars depict standard error of the mean. n = 3 and 4 for mean log_2_(FC) in gene expression and mean bacterial density, respectively.

### Genome-wide transcriptional changes indicate strong similarity in the plant response to various bacterial leaf colonisers

The strongest changes in gene expression were observed in Pst, Willi354 and Micro347 treated plants at 96 hpi. To gain an understanding of genome-wide transcriptional changes to individual leaf-colonising strains, the RNA of plants sampled at 96 hpi treated with either Pst, Willi354 or Micro347 were further subjected to RNA sequencing. The expression profiles of various target genes quantified by RNA sequencing were similar to those quantified by RT-qPCR confirming that RNA sequencing was performed correctly (Fig. **S2**). As expected, the transcriptomes of mock-treated plants were distinct from those of inoculated plants, as shown by multi-dimensional scaling (MDS) and k-means clustering (Fig. **4a**, Fig. **S3c**). The transcriptomes separate by treatment along the first dimension of the MDS plot (Fig. **4a**). Mock and Pst treated samples separate the furthest, corresponding to a leading FC of ∼ 4-fold between the two treatments. Micro347 and Willi354 treated samples cluster close to Pst with Willi354 being closer to Pst than Micro347, indicating a higher overlap of differentially expressed genes (DEGs). The second dimension of the MDS plot mainly separates the individual transcriptomes within a treatment group, showing low variation within Micro347 samples and strong variation within Willi354 samples, corresponding to a leading FC of ∼ 3-fold (Fig. **4a**).

**Fig. 4:**
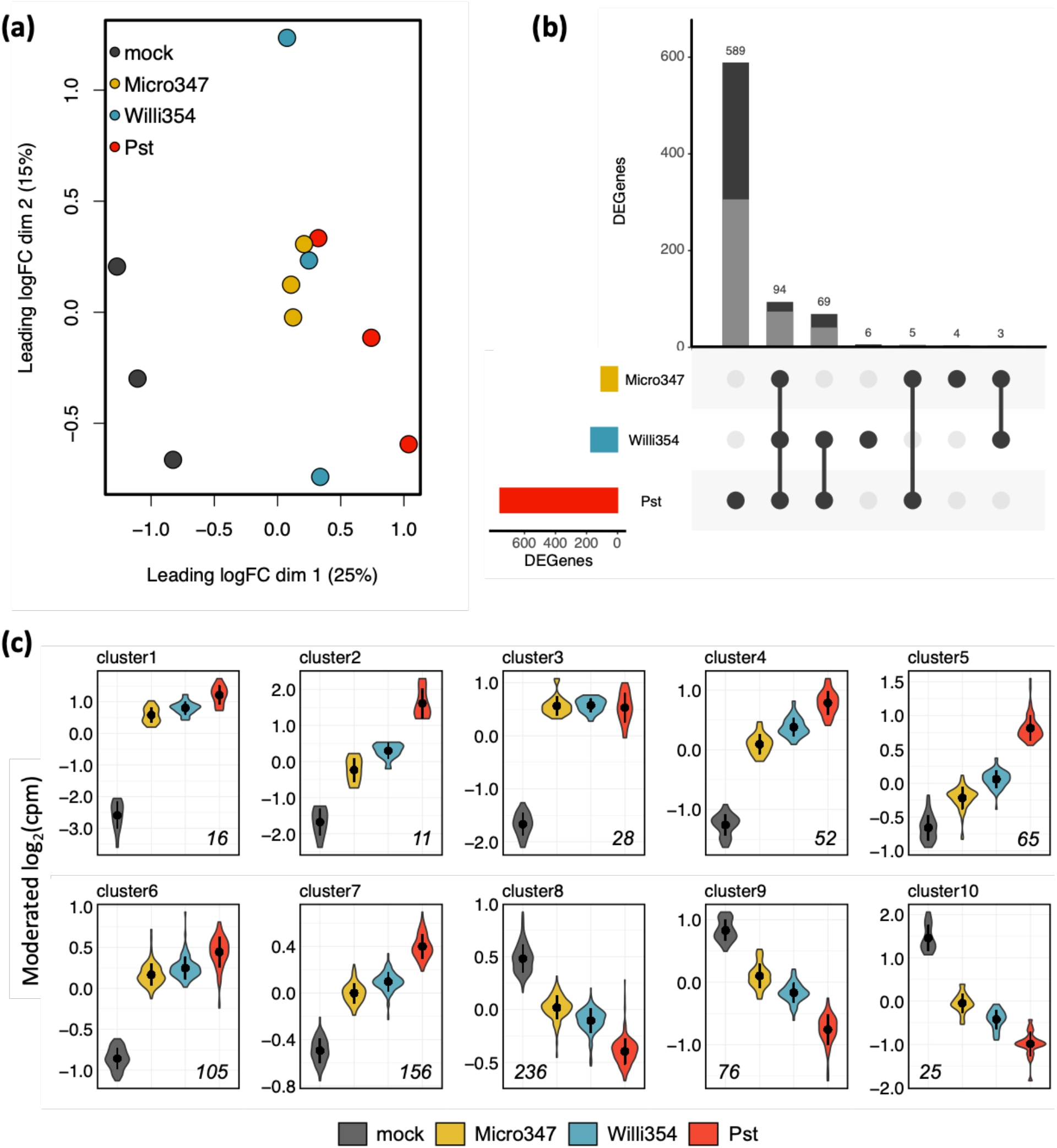
Comparison of transcriptomic response to various leaf-colonising strains. (**a**) Multidimensional scaling plot of transcriptomes of above ground plant parts of four-weeks-old axenically-grown arabidopsis plants four days after spray-inoculation with individual bacterial strains. Points depict individual transcriptomes, colour depicts inoculant or mock control treatment. (b) UpSet plot of DEGs in the different treatments. The coloured bar chart on the left depicts the total number of DEGs per treatment. The black dots in the panel’s matrix depict unique (individual dots) and overlapping (connected dots) DEGs. The top bar chart depicts DEGs for each unique or overlapping combination in the panel’s matrix. Grey bar depicts upregulated DEGs, black bar depicts downregulated DEGs. (**c**) Plots showing moderated log_2_(cpm) of DEGs per k-means cluster of transcriptomes of above ground plant parts of four-weeks-old axenically-grown arabidopsis plants four days after spray-inoculation with individual bacterial strains. Colours depict the treatment, the italicised number in each plot depicts the number of DEGs per cluster. Treatments were ordered based on the number of DEGs. Note that the y-axis differs between different cluster-plots.

Genes with a significant FC threshold of log_2_(1.3) based on the edgeR TREAT algorithm (McCarthy & Smyth, 2009) and a FDR < 0.05 were defined as DEGs. This FC threshold was chosen, based on the median ‘elbow’, the point of maximum curvature (second derivative) of the number of DEGs as a function of the log_2_(FC) threshold per treatment (Fig. **S3a**). Pst treated plants exhibited 757 DEGs at this FC threshold, followed by Willi354 with 172 DEGs and Micro347 with 106 DEGs. Interestingly, almost all the DEGs in Micro347 are also differentially expressed in Willi354 and Pst. In addition, almost all the DEGs found in Willi354 are also differentially expressed in Pst (Fig. **4b**). This shows that plant responses to these three leaf-colonising strains are largely similar but differ in their strength depending on the leaf coloniser. A closer look at individual gene expression changes further highlights the similarity in the responses. Most changes follow a sequence with either increasing or decreasing expression from mock-treated control over Micro347 and Willi354 to Pst-treated plants (Fig. **4c**, Fig. **S3c**). Further, all the 770 genes that were differentially expressed in any of the treatments, were either up- or downregulated in all the treatments. No gene was significantly upregulated in one treatment and significantly downregulated in another.

To gain a better resolution of gene expression changes, the 770 genes that were differentially expressed in any of the treatments were further separated by k-means clustering. K-means clustering was performed based on moderated log_2_(cpm). Ten k-means were chosen, based on the ‘elbow’ of the total sum of squares as a function of the number of k-means (Fig. **4c**, Fig. **S3b,c**). The 433 upregulated genes are in clusters 1-7, and the 337 downregulated genes are in clusters 8-10 (Fig. **4c**, Fig. **S3c**). In addition to more genes being significantly upregulated than downregulated, FCs were greater in upregulated genes. Clusters 1 and 2 contain genes with the strongest upregulation and cluster 10 genes with the strongest downregulation at FCs in moderated log_2_(cpm) of ∼ 16-fold and ∼ 6-fold, respectively (Fig. **4c**).

### Transcriptional responses depend on bacterial load

As seen above, responses to bacterial colonisation seem to be largely similar (Fig. **4b,c**) in response to the tested strains. This was especially surprising as Pst is an arabidopsis pathogen, whereas Micro347 and Willi354 were isolated from leaves of asymptomatic plants (Cuppels, 1986; Bai *et al*., 2015). In addition, bacterial density, irrespective of the bacterial coloniser, had a highly significant effect on ethylene responses (Fig. **3**). Taken together, this suggests that pathogenicity is to some extent dependent on bacterial density, following Paracelcus’ theory “the dose makes the poison” (Paracelsus, 1538). This raises the question whether non-pathogenicity of bacteria is merely a case of the plant balancing their proliferation, or the bacteria doing so in order to avoid being penalised by the plant.

To gain a better understanding of plant responses to non-pathogenic leaf-colonising bacteria and to determine whether the bacterial load changes the nature of the response, plants were inoculated with different concentrations of Willi354 followed by RNA sequencing. Willi354 was chosen for this experiment as it exhibited stronger responses than Micro347 in the previous experiment (Fig. **4b,c**).

Six-weeks-old axenically-grown arabidopsis plants were spray inoculated with Willi354 with inoculation densities ranging from 10^5^ CFU ml^-1^ to 10^8^ CFU ml^-1^. Four days after inoculation, the bacterial densities on the plants ranged from 6.57 × 10^6^ CFU g^-1^ to 3.22 × 10^8^ CFU g^-1^ and strongly correlated with the inoculation density (adj.R^2^ = 0.9265, *p* = 3.594 × 10^-14^) (Fig. **5a**). The transcriptomic response of the plants changed gradually with increasing inoculation density, as seen in the MDS plot and k-means clustering (Fig. **5b**, Fig. **S4**). The transcriptomes separate along the first dimension of the MDS plot, which explains 55% of the variation between the different samples, with those of mock-treated plants and those of plants inoculated with Willi354 at 10^8^ CFU ml^-1^ being most dissimilar at a leading FC of ∼ 16-fold (Fig. **5b**).

**Fig. 5:**
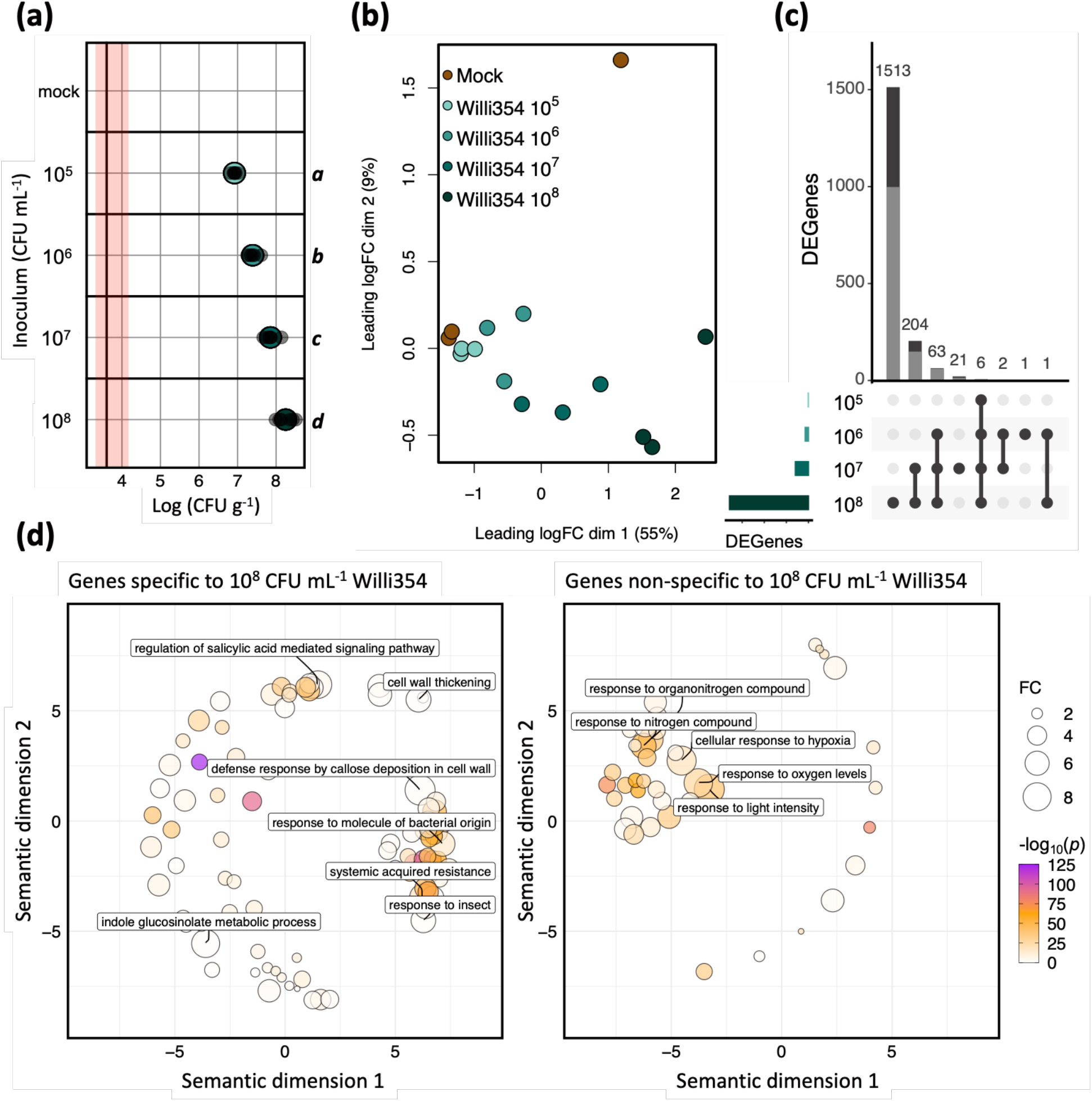
Plant transcriptomic response to different densities of Willi354. (**a**) Bacterial density of Willi354 on above ground plant parts of six-weeks-old axenically-grown arabidopsis plants 96 hpi with different initial densities. Coloured circles depict the mean, bars depict standard deviation, smaller grey circles depict the bacterial density from individual biological replicates, the vertical black line represents the threshold of detection based on mean plant weight, red bar represents the threshold range of detection based on the heaviest and lightest plant. Letters on the right side of the plot depict statistical differences (*p* < 0.001, one-way ANOVA & Tukey’s HSD test). (**b**). Multidimensional scaling plot of transcriptomes of arabidopsis above ground plant parts 96 hpi. Circles depict individual transcriptomes, colour depicts inoculation density or mock control treatment. (**c**) UpSet plot of transcriptomes depicted in (**b**). Genes with a FC significantly above log_2_(1.3) and a FDR < 0.05 were defined as DEGs. In each panel, the bottom left bar chart depicts the overall number of DEGs per treatment. The dots in the panel’s matrix depict unique (individual dots) and overlapping (connected dots) DEGs. The top bar chart depicts DEGs for each unique or overlapping combination in the panel’s matrix. Grey bar depicts upregulated DEGs, black bar depicts downregulated DEGs. (**d**) Functional enrichment of genes specifically and non-specifically expressed to high densities of Willi354. Functional enrichment of GO terms distributed in the semantic space. Closeness in semantic space ideally reflects closeness in GO term structure. Circles depicted significantly enriched GO terms (*p* < 0.05, Bonferroni corrected). Circle size depicts fold enrichment, colour depicts -log_10_(*p*). GO terms with a FC > 6 are labelled.

Genes were defined as DEGs as described above. Plants treated with 10^5^, 10^6^, 10^7^ and 10^8^ CFU ml^-1^ of Willi354 exhibited 6, 73, 296 and 1787 DEGs, respectively. A total of 1811 genes were differentially expressed across all treatments, with ∼ 68% (1230 DEGs) being upregulated (Fig. **5c**). Further, upregulated genes exhibited stronger changes in gene expression, the strongest being ∼ 30-fold in moderated log_2_(cpm), compared to downregulated genes, the strongest being ∼ 4-fold in moderated log_2_(cpm) (Fig. **S5**). This highlights that positive expression changes are not only more prominent, but also stronger than negative expression changes. The DEGs were separated by k-means clustering into 10 clusters.

Clusters 1-8 contained the upregulated, and clusters 9 and 10 contained the downregulated genes (Fig. **S4c**, Fig. **S5**). Most genes showed no marked difference between mock-treated plants and plants treated with Willi354 at 10^5^ and 10^6^ CFU ml^-1^. At concentrations of 10^7^ and 10^8^ CFU ml^-1^, genes in clusters 1, 3, 4,6 and 8 were strongly upregulated, with genes at 10^8^ being upregulated twice as much compared to those at 10^7^ (Fig. **S5**). Genes in clusters 5 and 7 appear to follow a sigmoidal curve, with no difference in gene expression between plants inoculated with Willi354 at 10^7^ and 10^8^ CFU ml^-1^ in cluster 7 (Fig. **S5**). Genes in clusters 9 and 10 exponentially decreased in expression with increasing density of Willi354 (Fig. **S5**).

To further explore the nature of plant responses to inoculation with Willi354, a GO term enrichment analysis was performed on two different sets of genes. The first set comprised genes that were exclusively differentially expressed in plants treated with Willi354 at 10^8^ CFU ml^-1^, whereas the second set comprised genes that were also differentially expressed at lower densities of Willi354. The functional profiles of both sets of genes were markedly different (Fig. **5d**). Genes differentially expressed exclusively at the highest density of Willi354, were greatly enriched for genes related to plant immunity, including perception of the biotic environment, such as ‘response to molecule of bacterial origin’ and ‘response to insect’, metabolism of secondary metabolites such as ‘indole glucosinolate metabolic process’, local defence responses such as ‘cell wall thickening’ and ‘defence response by callose deposition in cell wall’, and systemic defence responses such as ‘regulation of salicylic acid mediated signalling pathway’ and ‘systemic acquired resistance’ (Fig. **5d**). By contrast, genes that were also differentially expressed at lower densities of Willi354, were greatly enriched for genes related to plant nitrogen homeostasis, such as ‘response to nitrogen compound’ and ‘response to organonitrogen compound’, plant oxygen levels, such as ‘response to oxygen levels’ and ‘response to hypoxia’ and the plant’s ‘response to light intensity’ (Fig. **5d**).

### Transcriptional changes induced by bacteria may depend on the chromatin state of a given gene

A major determinant of gene transcriptional regulation is the chromatin state of genes. Chromatin states are determined by the combination of chromatin modifications and histone variants. Various histone modifications, such as H3K4me3, H3K36me3 and lysine acetylation, were previously shown to contribute to the induction of genes in response to pathogen exposure (Berr *et al*., 2012; Ding & Wang, 2015). As genes specifically induced by high densities of Willi354 presented distinct functional enrichment than genes induced by Willi354 irrespective of inoculation density, we hypothesised that both of sets of genes, genes activated after high inoculation and genes activated irrespectively of inoculation density, might exhibit different chromatin states prior to inoculation. To that end, the state of both sets of genes was examined using the chromatin state topology established by Sequeira-Mendes et al. (2014) (Fig. **6a**). Both sets exhibited an enrichment of states 1 and 2, which are both characterised by the presence of the histone variant H2A.Z accompanied either by activating marks such as H3K4me3, H3K36me3 or by a combination of activating (H3K4me3) and repressive (H3K27me3) marks, respectively. They also displayed an underrepresentation of states 8 and 9, which contain heterochromatic marks such as H3K9me2 and H3K27me1. In addition, genes which are upregulated by Willi354 irrespective of inoculation density showed an underrepresentation of states 3, 5, 6 and 7. By contrast, genes solely induced by the highest density of Willi354 either showed an enrichment (state 6) or no difference to the reference. These results reveal that, similarly to the distinct functional enrichments, genes induced specifically by higher bacterial densities exhibit a partially different chromatin signature than genes that are also induced by lower densities.

**Fig. 6:**
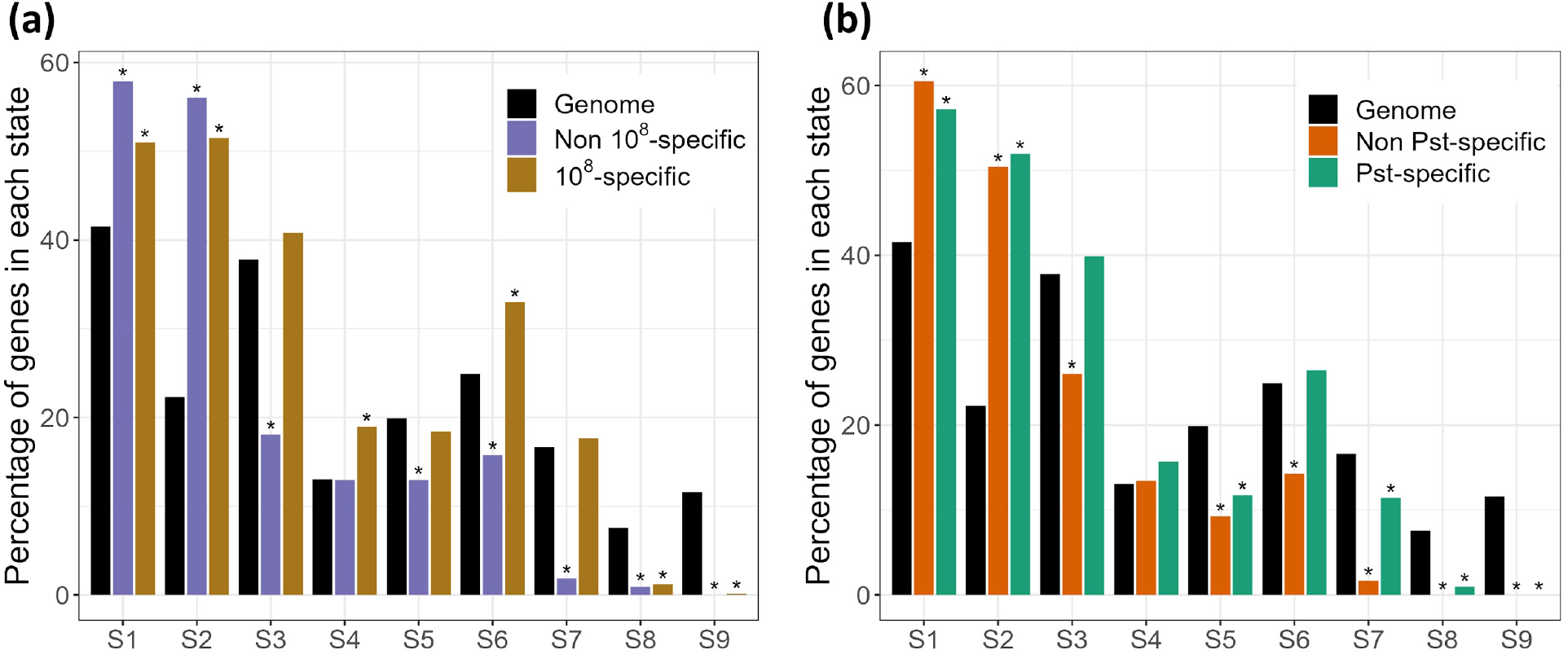
Chromatin state analysis of genes upregulated following bacterial inoculation. (**a**). Chromatin state analysis of genes induced either specifically by the highest density (10^8^) of Willi354 or also induced by lower inoculant densities (10^5^, 10^6^ or 10^7^). (**b**) Chromatin state analysis of genes induced either specifically by Pst or also induced by Willi354 or Micro347 (all at a density of 10^7^). The chromatin states coordinates were obtained from Sequeira-Mendes et al. (2014). * indicates significant difference compared to the genome, tested by Marascuilo procedure (α = 0.05).

As the genes induced solely by the higher densities of Willi354 displayed an enrichment for plant immunity and defence-related terms (Fig. **5d**), we wondered whether those genes would have a similar chromatin profile as genes induced by Pst. The genes were divided into two sets, depending on whether they were induced specifically by Pst or whether they were also induced by Willi354 or Micro347 (Fig. **6b**). Similarly, to the previous analysis, both sets presented an enrichment for states 1 and 2 and an underrepresentation for states 8 and 9. They also showed an underrepresentation of state 5, characterised by the presence of the repressive mark H3K27me3. Additionally, genes induced by both Pst or Willi354/Micro347 displayed an underrepresentation of states 6 and 7, which was not observed or of reduced magnitude for the genes induced specifically by Pst. Despite limited overlap between the two transcriptomic experiments (Fig. **S6**), the chromatin profiles of genes induced specifically either by high Willi354 densities or Pst were comparable and distinct from the profiles of genes induced also by non-pathogenic bacteria or by lower bacterial densities. This observation supports the idea that inoculation with higher densities of non-pathogenic bacteria leads to a transcriptomic response similar to that triggered by pathogenic bacteria.

### High densities of Willi354 caused slight disease phenotypes

Since the genes that were uniquely differentially expressed in plants inoculated with Willi354 at 10^8^ CFU ml^-1^ were enriched for plant immunity-related genes, plants were inoculated with Willi354 under the previous experimental conditions and sampled at 14 and 21 dpi, to investigate if Willi354 evokes plant disease phenotypes. Indeed, plants inoculated with Willi354 at 10^8^ CFU ml^-1^, but not at lower densities, exhibited necrotic lesions on a few leaves 14 and 21 dpi (Fig .**7a-f**, Fig. **S7**). In addition, plant weight was negatively correlated with inoculation density of Willi354 (adj.R^2^ = 0.11, *p* = 6.48 × 10^-5^) (Fig. **7h**).

**Fig. 7:**
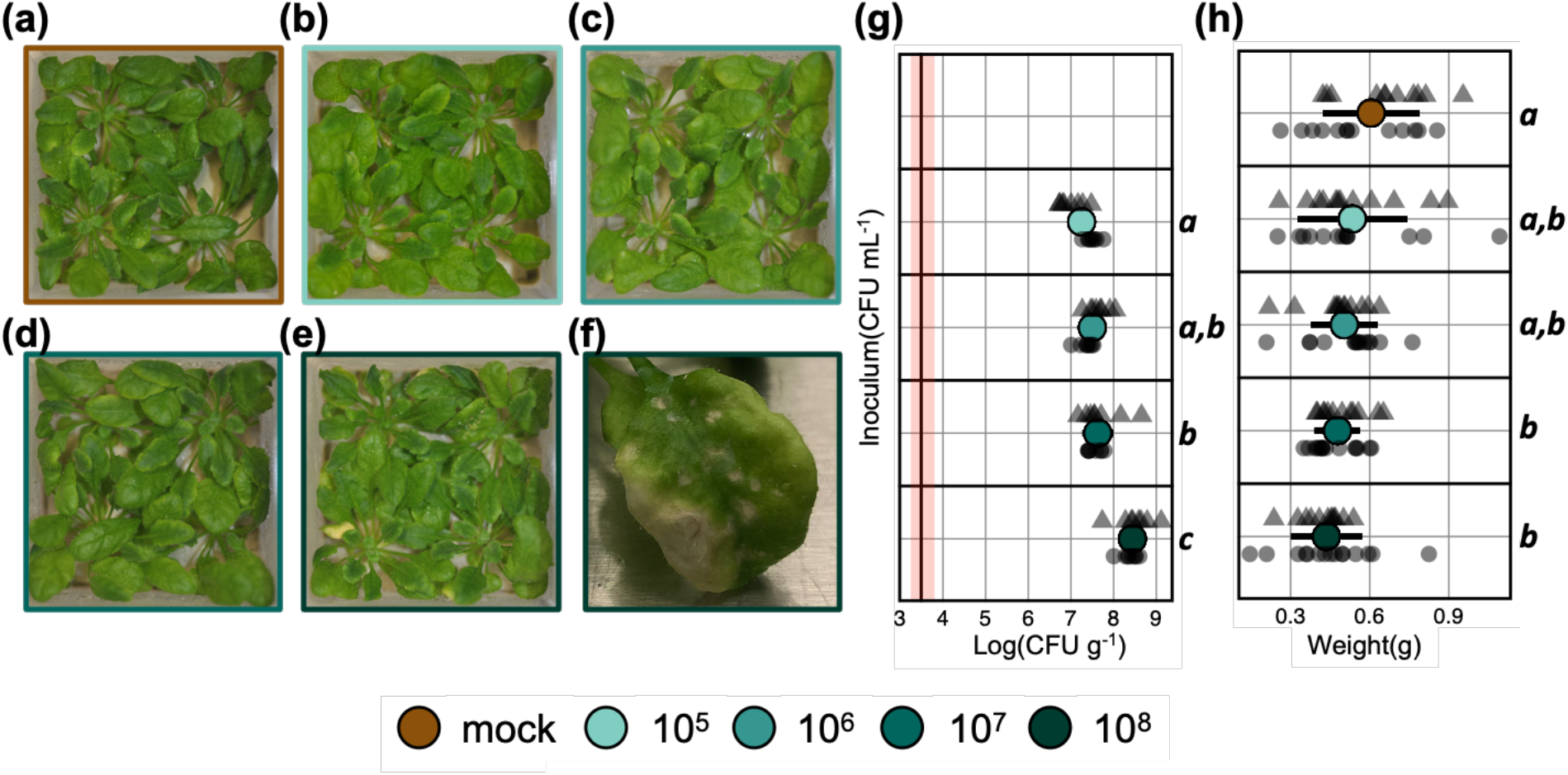
Effect of Willi354 density on plant phenotype 21 dpi. Representative images of six-weeks-old axenically-grown arabidopsis plants. Plants were either mock-treated (**a**) or inoculated with Willi354 at 10^5^ CFU ml^-1^ (**b**), 10^6^ CFU ml^-1^ (**c**), 10^7^ CFU ml^-1^ (**d**) or 10^8^ CFU ml^-1^ (**e**). (**f**) Representative image of the leaves that showed spots of localised cell death in plants inoculated with Willi354 at 10^8^ CFU ml^-1^. (**g**) Bacterial density of Willi354 on aboveground plant parts of six-weeks-old axenically-grown arabidopsis plants. The vertical black line represents the threshold of detection based on mean plant weight, the red bar represents the threshold range of detection based on the heaviest and lightest plant. (**h)** Fresh weight of six-weeks-old axenically-grown arabidopsis plants. Coloured circles depict the mean, bars depict standard deviation, smaller grey shapes depict the bacterial density from individual biological replicates of two independent experiments. Letters on the right side of the plots depict statistical differences (*p* < 0.05), one-way ANOVA & Tukey’s HSD test.

## Discussion

### Temporal responses of the plant immune system

Bacteria were spray-inoculated at densities that matched the bacterial carrying capacity of leaves in temperate environments (Kniskern *et al*., 2007; Reisberg *et al*., 2012; Rastogi *et al*., 2012; Burch *et al*., 2016; Gekenidis *et al*., 2017). Four of the six inoculated bacteria successfully established on plant leaves, whereas two, Acido84 and Pedo194, failed to consistently reach bacterial densities above the threshold of detection, which was on average ∼ 2500 CFU g^-1^ of leaf fresh weight (Fig. **2a**). This was rather surprising as both genera were previously found to make up more than 1% of the total bacterial population on arabidopsis (Vorholt, 2012). Further, both strains were recently shown to successfully colonise arabidopsis (Vogel *et al*., 2021). However, in the study by Vogel and colleagues, colonisation density was measured nine days after drop inoculation on seedlings in an agar-based system. In this study the ‘Litterbox’ system was employed, which reliably mimics environmental population densities, as opposed to agar-based systems that exhibit unnaturally high population densities (Miebach *et al*., 2020). In addition, Acido84 reached population densities of 10^4^ - 10^6^ CFU g^-1^ in all sampled plants at 168 hpi (i.e. 7 dpi). Since Acido84 was also successfully recovered straight after inoculation this indicates (1) that Acido84 was not harmed during the spraying procedure and (2) that it was able to thrive on leaves at later time points. Overall, this suggests that Acido84 had to acclimatise to its new environment after growth on R2A media, even though R2A is, like the phyllosphere, oligotrophic. Pedo194, in contrast, was only successfully recovered from one plant immediately after spray-inoculation at a density ∼ 2 magnitudes lower than the inoculum, suggesting that the inoculation procedure might have been detrimental to it.

Pst and Will354 rose in population size to ∼ 10^7^ CFU g^-1^ at 96 hpi. Population size then declined to 10^6^ CFU g^-1^ at 168 hpi (Fig. **2a**). Whether this was due to exhaustion of resources or plant immune responses remains to be determined. Interestingly though, the expression of the ET marker *ARL2* could be explained in large parts by the bacterial density of the coloniser, irrespective of the inoculant. This suggests that *ARL2* expression must be either triggered by a common MAMP, shared between the isolates, or was triggered by many MAMPs which the plant did not distinguish between. Further, it suggests that *ARL2* expression is proportional to the MAMP titer. This agrees with previous findings, which described stronger transcriptional responses to both higher pathogen and higher MAMP titers (Thilmony *et al*., 2006; Denoux *et al*., 2008).

The observed expression changes upon bacterial treatment were overall rather weak (Fig. **2b**). The strongest changes in gene expression were observed in the ET marker, *ARL2*, and culminated at 96 hpi. As expected, the strongest changes were observed in the plants treated with Pst. The overall weak and rather late response seemingly disagrees with previous studies describing fast and substantial changes in gene expression upon MAMP treatment and infection with Pst (Thilmony *et al*., 2006; Zipfel *et al*., 2006; Denoux *et al*., 2008; Bjornson *et al*., 2021). However, in these studies either young seedlings were treated by a complete change of media, with the fresh media containing the MAMP or leaves of mature plants were vacuum infiltrated. In both cases MAMPs were readily available. By contrast, in the case of a surface spray, a sufficient amount of eliciting molecules needs to cross the hydrophobic cuticle layer to reach the plasma membranes of plant cells (Schlechter *et al*., 2019) or bacteria need to migrate into the apoplast (Beattie & Lindow, 1999; Melotto *et al*., 2006). In addition, the flg22 receptor, FLS2 is highly expressed in leaves near bacterial entry sites, such as stomata, which are predominantly found on the abaxial (lower) leaf surface and hydathodes, as well as in leaf veins (Beck *et al*., 2014). Vacuum infiltration of bacterial suspensions would render leaf veins more exposed to MAMPs and a change in liquid media would render stomata and hydathodes more exposed to MAMPs, than in the more ‘natural’ scenario of topical application.

### Genome-wide transcriptional responses to leaf colonisation

#### Plant responses to non-pathogenic bacteria are qualitatively similar but differ quantitatively compared to pathogenic bacteria

Bacterial colonisation with non-pathogenic leaf colonisers significantly altered the expression of several genes in the host, although not to the extent of a pathogenic leaf coloniser. Remarkably, the responses observed were largely similar, but weaker in response to colonisation by the non-pathogenic bacteria. Most genes significantly expressed in response to one strain were also significantly expressed in response to strains that elicited stronger responses and thus, higher numbers of DEGs. None of the 770 genes with a significant FC in any of the bacterial treatments was upregulated by one strain and downregulated by another. Changes in gene expression of genes belonging to clusters 2, 4, 5, 6, 7, 8, 9 and 10 were either progressively increasing or decreasing when treatments were sorted by the number of DEGs that they elicited (Fig. **4c**). This indicates that those genes were similarly regulated in response to bacterial colonisation irrespective of the symbiotic relationship of the inoculant with the plant, although less severely in response to non-pathogenic bacteria. Such similarity in the plant response to various leaf colonisers was also described recently by Maier and colleagues, although without the context of a pathogenic bacterium (Maier *et al*., 2021). Interestingly, the response strength was strongly driven by the bacterial density of the inoculant (Maier *et al*., 2021), which was also observed in the current study with respect to ET marker responses to non-pathogenic and pathogenic bacteria. This suggests that plants merely responded to a pool of bacterial MAMPs quantitatively, by responding to the total amount of MAMPs present, rather than qualitatively by integrating a unique mix of different MAMPs into a tailored plant response.

### The effect of bacterial load on plant gene expression

Bacterial densities were the major driver of ethylene marker expression between 24 and 96 hpi irrespective of the bacterial coloniser (Fig. **3**). In addition, genome-wide transcriptional responses to bacterial colonisation were largely similar but differed in number of DEGs (Fig. **4b**), as well as in the expression strength of individual genes (Fig. **4c**). This was remarkable, as the tested strains included bacteria isolated from asymptomatic plants (Bai *et al*., 2015) as well as pathogenic Pst (Cuppels, 1986). As pathogenicity is linked to bacterial density, this raises the question whether non-pathogenicity is merely a case of the plant limiting uncontrolled proliferation via Pattern Triggered Immunity or bacteria limiting their proliferation to avoid being penalised by the plant.

Plants were inoculated with Willi354 at various concentrations, resulting in different bacterial densities ranging from somewhat natural densities ∼ 10^6^ - 10^7^ CFU g^-1^ to artificially high densities ∼ 10^8^ - 10^9^ CFU g^-1^ within 96 hpi (Fig. **5a**). These differences in bacterial densities were still observed at 21 dpi (Fig. **7g**). The maximal bacterial load, referred to as the carrying capacity, strongly correlated with the inoculation density (Fig. **5a**), as was previously demonstrated on bean leaves (Wilson & Lindow, 1994; Remus-Emsermann *et al*., 2012). Interestingly, severe disease phenotypes were observed on a few leaves of plants colonised by Willi354 at ∼ 10^8^ - 10^9^ CFU g^-1^ 14 and 21 dpi (Fig. **7e,f**, Fig. **S7**). In addition, plant weight negatively correlated with bacterial density (Fig. 7g,h). This suggests that bacteria that are otherwise non-pathogenic can be detrimental to the plant at very high densities.

RNA sequencing revealed an exponential increase in both the number of DEGs and the expression pattern of most DEGs to increasing bacterial densities (Fig. **5c**, Fig. **S5**). In addition, gene expression changes were relatively strong in upregulated genes with FCs up to ∼ 30-fold (Fig. **S5**). This suggests that the plant barely invests energy into an interaction with its bacterial colonisers at low bacterial densities, but drastically increases responses when bacteria are reaching potentially dangerous levels. Accordingly, genes that were uniquely differentially expressed in plants harbouring Willi354 at ∼ 10^8^ - 10^9^ CFU g^-1^ were greatly enriched for immune related GO terms, including perception of the biotic environment (‘response to molecule of bacterial origin’, ‘response to insect’), metabolism of secondary metabolites (‘indole glucosinolate metabolic process’), local defence responses (‘cell wall thickening’, ‘defence response by callose deposition in cell wall’), and systemic defence responses (‘regulation of salicylic acid mediated signalling pathway’, ‘systemic acquired resistance’) (Fig. **7d**).

Because the plant ‘ignores’ bacteria at low densities, it appears unlikely that the plant limits bacterial proliferation by active signalling processes at low bacterial densities. Consequently, it is likely that bacteria limit their proliferation in order not to alert the plant immune system. This is in agreement with recent findings showing endophytic bacteria remaining in population stasis by a multiplication-death equilibrium independent of bacterial density (Velásquez *et al*., 2022).

The chromatin state analysis further supported the idea that artificially high densities of non-pathogenic bacteria lead to a transcriptomic response similar to that triggered by pathogenic bacteria. Genes induced by bacterial inoculation, both at pathogenic and non-pathogenic level, were enriched for the chromatin states 1 and 2 (Fig. **6**). In more detail they were enriched in H2A.Z and H3K4me3-H3K27me3 marks, previously shown to be essential for gene responsiveness to environmental changes (Coleman-Derr & Zilberman, 2012; Sura *et al*., 2017; Faivre & Schubert, 2023).This suggests that these chromatin marks might be guiding the transcriptomic response of the plant to bacterial inoculation.

## Conclusion

We show that plant responses to various leaf colonising bacteria are largely similar, both in the overlap of DEGs and the expression of individual DEGs but differ in expression strength. We tested the competing hypotheses that plants are either 1) monitoring bacterial population density or 2) differentiating between different bacterial colonisers. Our results suggest that plants are responding to bacterial densities rather than bacterial identities, favouring hypothesis 1) to be the more prevalent mode of plant response to bacteria. It appears that the plant barely invests resources into an interaction with its associated bacteria at low bacterial densities, but markedly increases gene expression towards defence responses upon high bacterial colonisation.

## Acknowledgements

This work was supported by Marsden Fast Start grant number 17-UOC-057 to M.N.P.R.-E. by the Royal Society Te Apārangi. M.M. was supported by a University of Canterbury Ph.D. scholarship.

## Author contributions

M.M., P.E.J. and M.N.P.R.-E. conceived and designed the study. M.M. performed all laboratory experiments and analysed the data. L.F. and D.S performed additional data analysis. M.M. wrote the initial draft of the manuscript and all authors contributed to later versions of the manuscript and to data interpretation.

