## Supplemental data for "Non-pathogenic leaf-colonising bacteria elicit pathogen-like responses in a colonisation density-dependent manner"

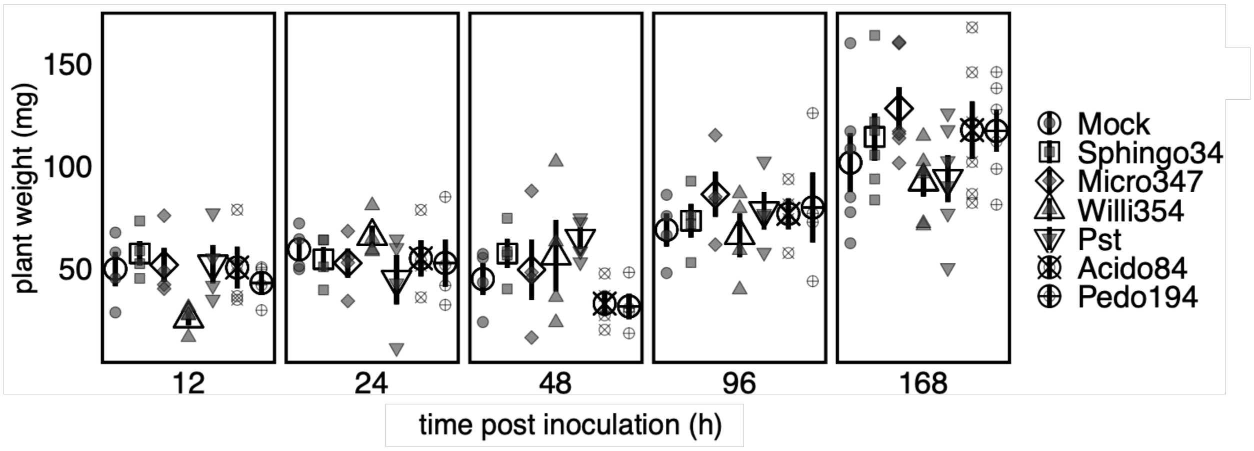


**Fig. S1: Plant weight post inoculation.** Four-weeks-old axenically-grown arabidopsis plants were spray-inoculated with individual bacterial strains. Bigger white shapes depict the mean, bars depict the standard error, smaller grey shapes depict the bacterial density from individual biological replicates, shapes represent the inoculant.

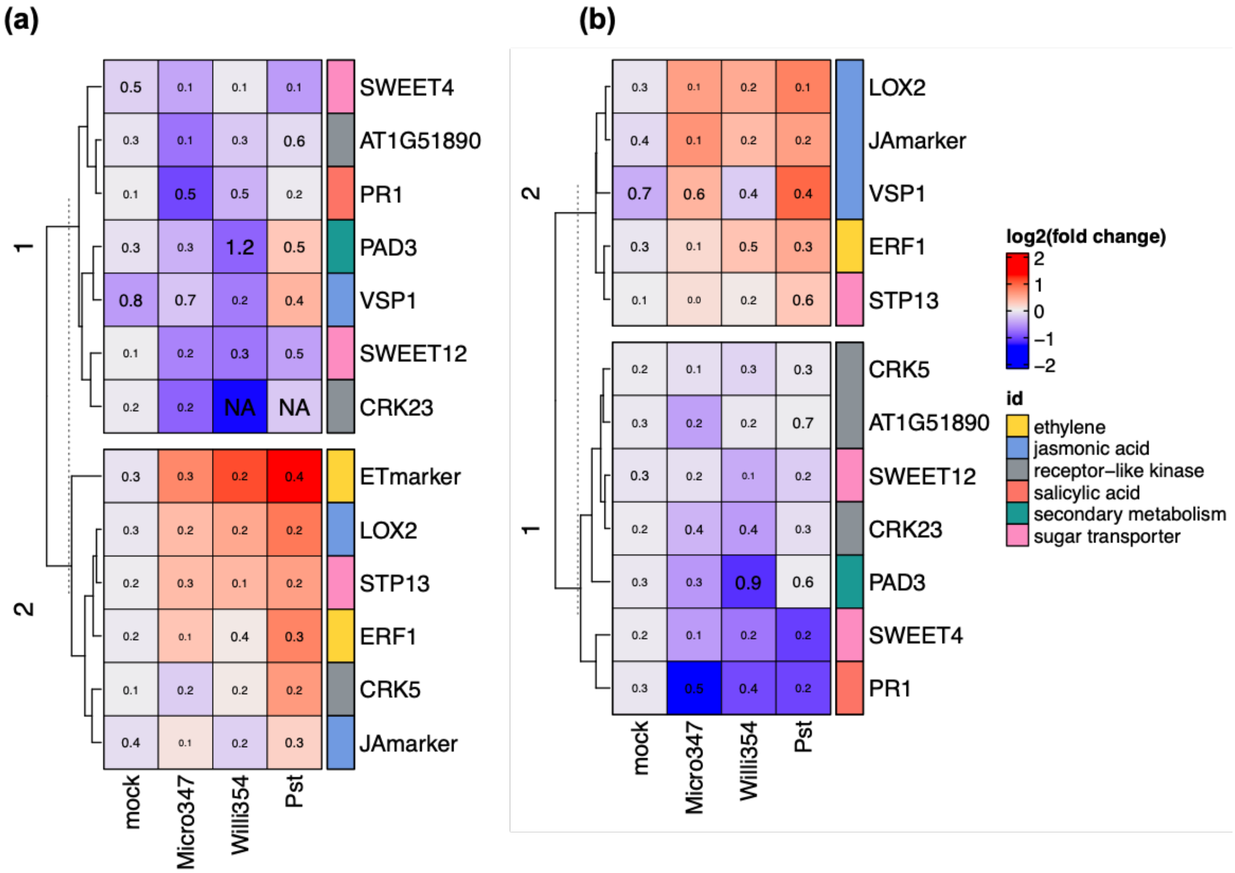


**Fig. S2: Selected genes differentially expressed to individual leaf colonisers.** Heatmaps showing $log_2$ FCs in transcripts of above ground plant parts of four-weeks-old axenically-grown arabidopsis plants four days after spray-inoculation with individual bacterial strains. (**a**) Heatmap of transcript changes determined via qPCR. (**b**) Heatmap of transcript changes determined via RNAseq. Gene names are shown on the right side of each plot and colour coded by affiliation to a functional category. Number inside each square represents the standard error.

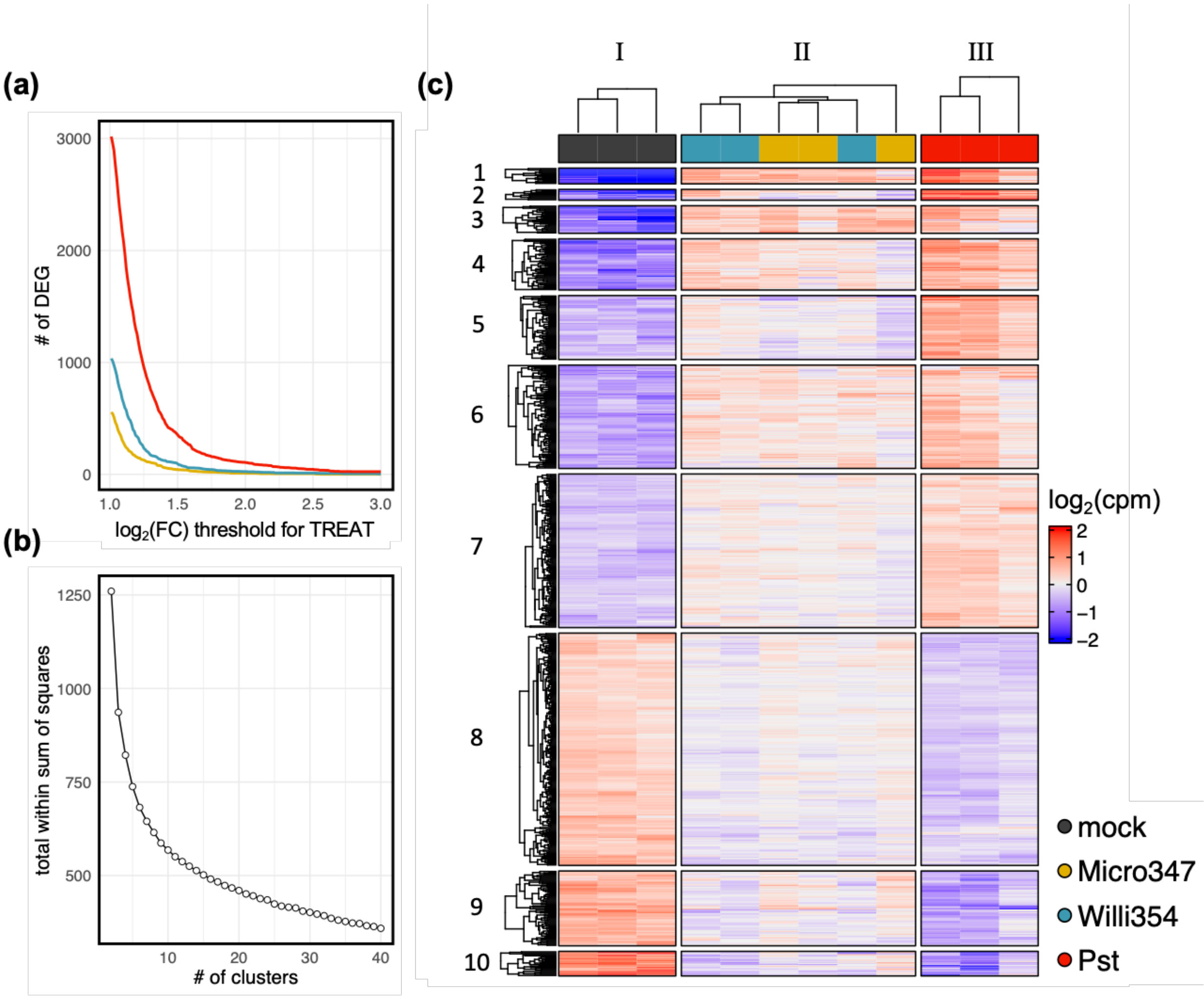


**Fig. S3: Gene expression response to various bacterial inoculants.** (**a**) Elbow plot showing the relation of FC threshold and the number of DEGs of transcriptomes of above ground plant parts of four-weeks-old axenically-grown arabidopsis plants four days after spray-inoculation with individual bacterial strains. Coloured lines depict the individual inoculants. (**b**) Elbow plot showing the relation of the total within sum of squares and the number of k-means clusters. (**c**) Heatmap of moderated $\log_2$(cpm) of DEGs. Colours on top of heatmap depict the different inoculants and the mock control treatment. Roman numbers on top and Arabic numbers on the left side of the plot depict the respective number of k-means clusters. Dendrograms represent hierarchical clustering based on euclidean distances for each k-means cluster.

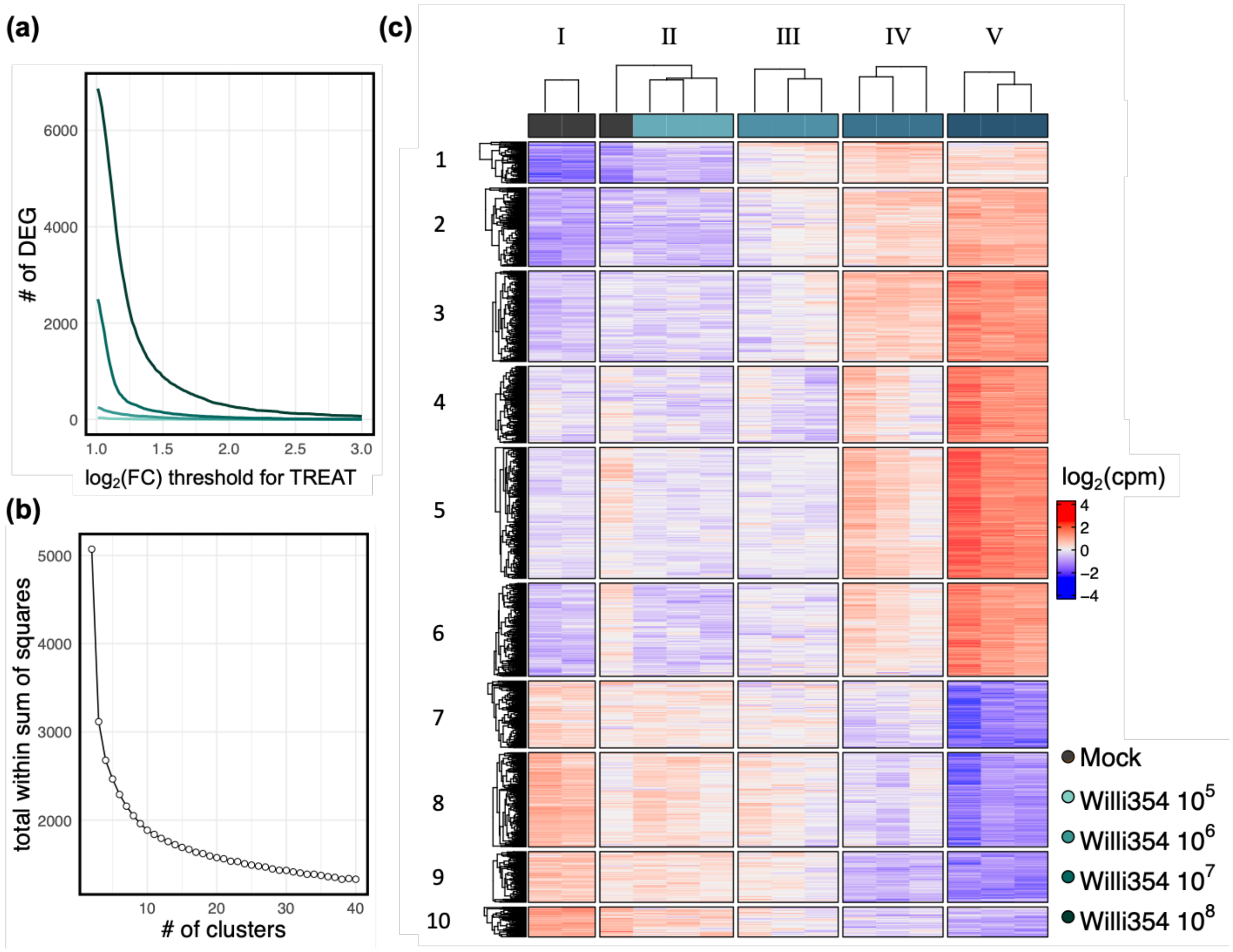


**Fig. S4: Gene expression response to different densities of Willi354.** (**a**) Elbow plot showing the relation of FC threshold and the number of DEGs of transcriptomes of above ground plant parts of six-weeks-old axenically-grown arabidopsis plants four days after spray-inoculation with Willi354 at different inoculum densities. Coloured lines depict the different inoculum densities. (**b**) Elbow plot showing the relation of the total within sum of squares and the number of k-means clusters. (**c**) Heatmap of moderated $\log_2$(cpm) of DEGs. Colours on top of heatmap depict the different inoculum densities and the mock control treatment. Roman numbers on top and Arabic numbers on the left side of the plot depict the respective number of k-means clusters. Dendrograms represent hierarchical clustering based on euclidean distances for each k-means cluster.

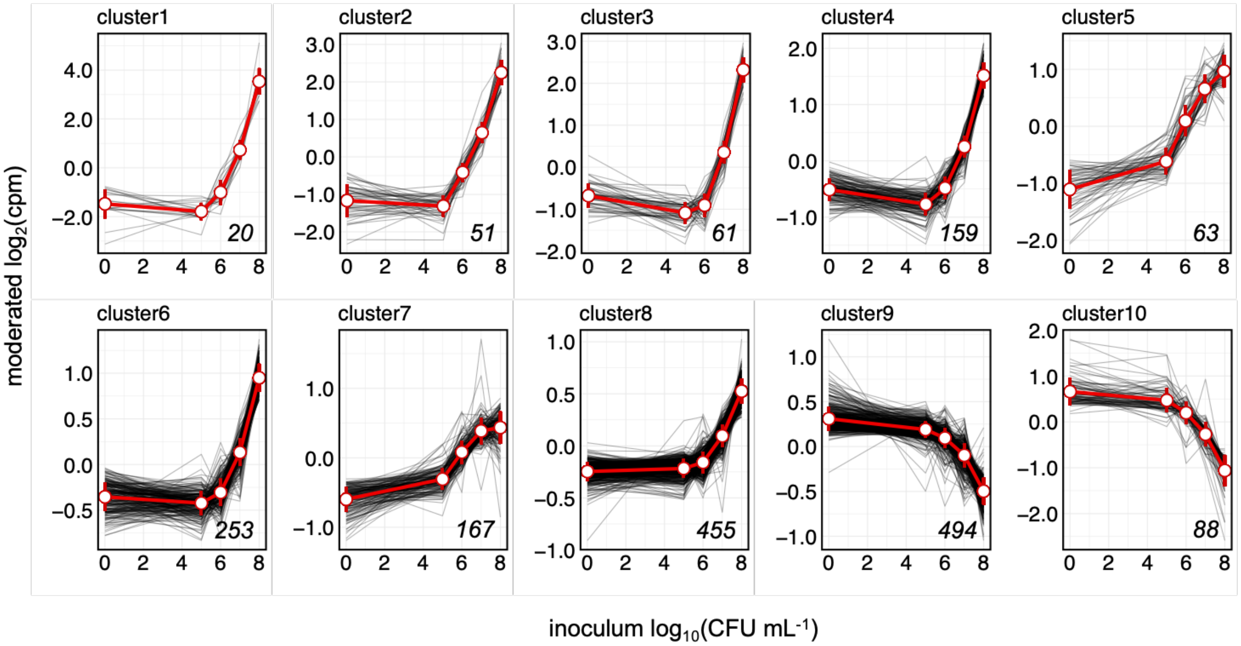


**Fig. S5: Individual gene expression changes to different densities of Willi354.** Plots showing moderated $\log_2$(cpm) of DEGs per k-means cluster of transcriptomes of above ground plant parts of six-weeks-old axenically-grown arabidopsis plants four days after spray-inoculation with different concentrations of Willi354. Red circles represent mean with standard deviation, grey lines represent individual genes, the italicised number in each plot depicts the number of DEGs per cluster. Note that the y-axis differs between different cluster-plots.

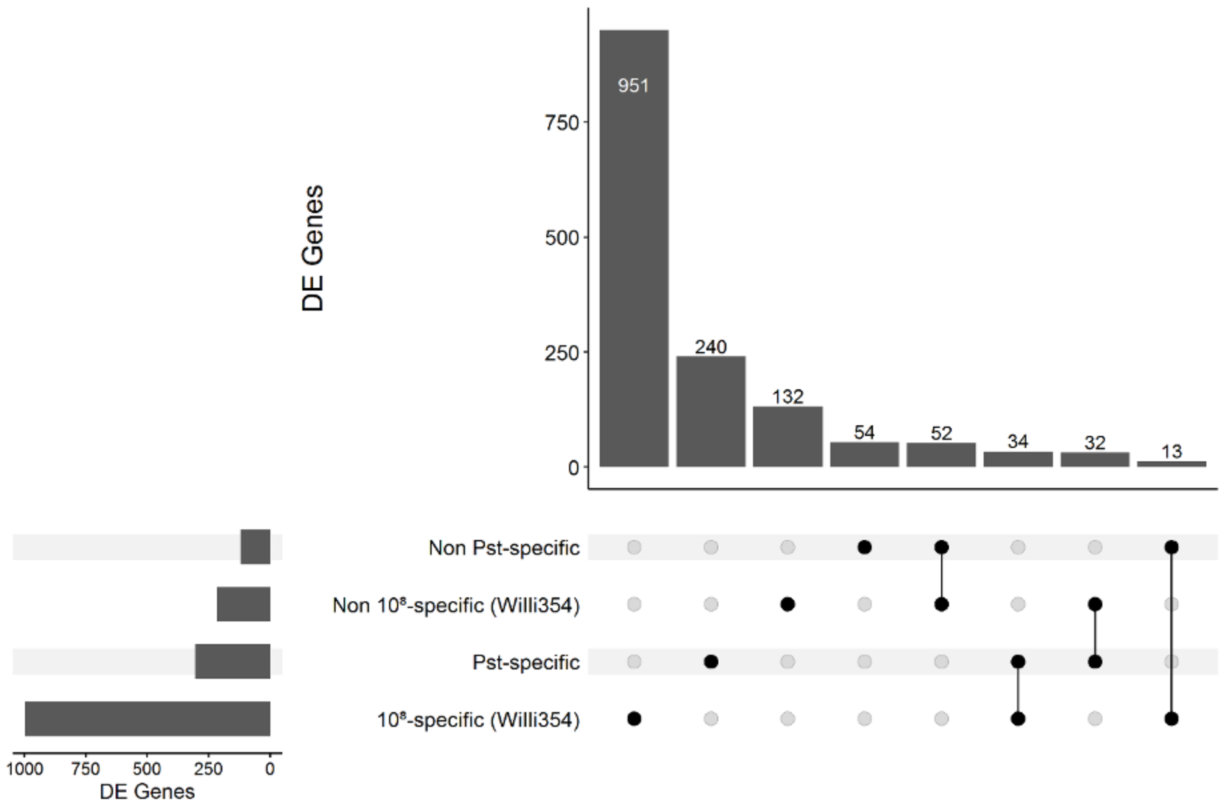


**Fig. S6: UpSet plot of the gene sets used in the chromatin state analysis.** The bar chart on the left depicts the total number of genes in each set. The black dots in the panel's matrix depict unique (individual dots) and overlapping (connected dots) genes. The top bar chart depicts genes for each unique or overlapping combination in the panel's matrix

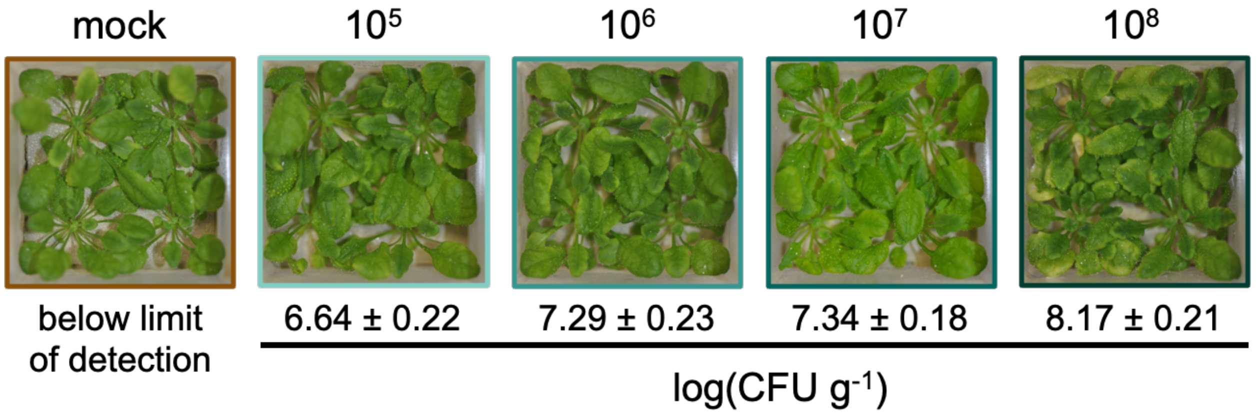


**Fig. S7: Effect of Willi354 density on plant phenotype 14 dpi.** Representative images of six-weeks-old axenically-grown arabidopsis plants. Plants were either mock-treated or inoculated with Willi354 at $10^5$, $10^6$, $10^7$ or $10^8$ CFU $ml^{-1}$, as represented by text above the images. Text below the images depicts the mean bacterial density and standard deviation of Willi354 on above ground plant parts of eight-weeks-old axenically-grown arabidopsis plants 14 dpi.
